# Genetically flexible but conserved: a new essential motif in the C-ter domain of HIV-1 group M integrases

**DOI:** 10.1101/2020.05.26.118158

**Authors:** Marine Kanja, Pierre Cappy, Nicolas Levy, Oyndamola Oladosu, Sylvie Schmidt, Paola Rossolillo, Flore Winter, Romain Gasser, Christiane Moog, Marc Ruff, Matteo Negroni, Daniela Lener

**Affiliations:** Université de Strasbourg, CNRS, Architecture et Réactivité de l’ARN, UPR9002, Strasbourg, France; Chromatin Stability and DNA mobility, Department of Structural Biology and Genomic, IGBMC, Strasbourg University, CNRS, INSERM, Illkirch, France; Molecular Immuno-Rhumatology Laboratory, UMR1109, FMTS, Université de Strasbourg, INSERM, Institut de Virologie, Strasbourg, France

**Author notes:** Corresponding authors: Daniela Lener and Matteo Negroni, E-mailing addresses. Mailing address: Institut de Biologie Moléculaire et Cellulaire, 2 allée Konrad Roentgen, 67084 Strasbourg Cedex, France.

## Abstract

Using coevolution-network interference based on the comparison of two phylogenetically distantly related isolates, one from the main group M and the other from the minor group O of HIV-1, we identify, in the C-terminal domain (CTD) of integrase, a new functional motif constituted by four non-contiguous amino acids (N_222_K_240_N_254_K_273_). Mutating the lysines abolishes integration through decreased 3’-processing and inefficient nuclear import of reverse transcribed genomes. Solution of the crystal structures of wt and mutated CTDs shows that the motif generates a positive surface potential that is important for integration. The number of charges in the motif appears more crucial than their position within the motif. Indeed, the positions of the K could be permutated or additional K could be inserted in the motif, generally without affecting integration *per se*. Despite this potential genetic flexibility, the NKNK arrangement is strictly conserved in natural sequences, indicative of an effective purifying selection exerted at steps other than integration. Accordingly, reverse transcription was reduced even in the mutants that retained wt integration levels, indicating that specifically the wt sequence is optimal for carrying out the multiple functions integrase exerts. We propose that the existence of several amino acids arrangements within the motif, with comparable efficiencies of integration *per se*, might have constituted an asset for the acquisition of additional functions during viral evolution.

**IMPORTANCE:** Intensive studies on HIV-1 have revealed its extraordinary ability to adapt to environmental and immunological challenges, an ability that is also at the basis of antiviral treatments escape. Here, by deconvoluting the different roles of the viral integrase in the various steps of the infectious cycle, we report how the existence of alternative equally efficient structural arrangements for carrying out one function opens on the possibility of adapting to the optimisation of further functionalities exerted by the same protein. Such property provides an asset to increase the efficiency of the infectious process. On the other hand, though, the identification of this new motif provides a potential target for interfering simultaneously with multiple functions of the protein.

## Introduction

Integration of reverse transcribed viral genomes into the genome of the infected cell is a peculiar feature of the replication strategy of retroviruses, carried out by the viral enzyme integrase (IN) in a two-step reaction. In HIV-1, after the achievement of DNA synthesis in the cytoplasm of the infected cell, it first catalyses the removal of a conserved GT dinucleotide from the 3’ ends of the viral DNA (3’ processing), leaving CA_-OH_ 3’ ends bound to the active site. Subsequently, once the viral DNA has been imported in the nucleus, the reactive CA_-OH_-3’ ends attack the cellular DNA leading to the generation of the provirus (1, 2).

Besides this enzymatic function, HIV-1 IN is involved, through non catalytic activities, in several other steps of the viral replication cycle. As a component of the Gag-Pol polyprotein precursor, it participates in Gag-Pol dimerization, essential for the auto-activation of the viral protease and, consequently, for viral particle maturation (3-5). During capsid morphogenesis, it is involved in the recruitment of the genomic RNA inside the core of the viral particle (6). As a mature protein, it interacts with the viral polymerase (reverse transcriptase, RT) to optimize reverse transcription of the viral genome (7-9). Through the interaction with the cellular protein LEDGF/p75, it targets actively transcribed genes as sites for integration (10). Finally, as a component of the pre-integration complex (PIC), IN is also involved in nuclear import of the reverse transcription product, a peculiar feature of lentiviruses that allows the infection of non-dividing cells.

This ability relies on the virus capacity to enter the nucleus *via* an active passage through the nuclear pore complex (NPC) (11, 12). Several lines of evidence have indicated that the capsid (CA) protein is crucial for nuclear entry (13, 14), through its interaction with several nucleoporins (Nups) forming the NPC (Nup 358, Nup 153, Nup 98) (15-17) and with the transportin-3 (18, 19). Nevertheless, several studies have indicated that IN has karyophilic properties. Namely, it contains a basic bipartite nuclear localization signal (NLS) (20) as well as an atypical NLS (21), and also binds several cellular nuclear import factors. Interactions of IN with importin α/β (22), importin 7 (23), importin α3 (24), Nup153 (25), Nup62 (26) and transportin-3 (27-29) have been documented. Indeed, the mutation of amino acids, mostly located in the C-terminal domain of the IN, responsible for binding to nuclear import factors, results in non-infectious viruses impaired in nuclear import (23, 24, 28, 30).

The functional form of the HIV-1 integrase is made up of a dimer of dimers, which assemble in highly ordered multimers of these tetramers (31, 32). Three domains, connected by flexible linkers, constitute HIV-1 IN: the N-terminal domain (NTD), the catalytic core domain (CCD) and the C-terminal domain (CTD) (33, 34). While the NTD is mostly involved in protein multimerization (35, 36), the CCD is mostly responsible for catalysis, and for binding to the viral and cellular DNA as well as to the cellular cofactor LEDGF (37-40). Finally, the CTD is involved in DNA binding during integration (41, 42), in protein multimerization (35), in the interaction with the reverse transcriptase (7, 9) and in the recruitment of the viral genomic RNA (gRNA) in the viral core (6). Overall, the intrinsic flexibility of the protein, the multiple steps required to achieve integration, and the multimeric nature of the integration complex make the involvement of the different parts of the protein in the various functions of the integrase very complex and still not fully elucidated.

In addition, the multiple tasks that the IN must accomplish during the infectious cycle and the complexity of its supramolecular structures are expected to impose functional constraints that ultimately may limit its genetic diversity. Retention of functionality despite sequence variation strongly relies on covariation, inside or outside the mutated protein. When an initial mutation negatively alters the protein functionality, compensatory mutations can restore it, at least partially. Therefore, the sequences of homologous proteins in different HIV variants are the result of independent evolution pathways, with independent covariation networks specifically generated for each pathway. Chimeric genes between variants of a given protein can perturb such networks and result in the production of non-functional proteins. This information can then be exploited to probe the existence of functional motifs in proteins. For considerably divergent viruses, as those derived from independent zoonotic transmissions, this approach can be particularly powerful. This is the case for HIV-1 groups M and O that derive from simian viruses infecting chimpanzees and gorillas, respectively. Here, we exploit the natural genetic diversity existing between these groups to generate chimeric integrases. A detailed characterization of the individual amino acids that differ in the non-functional chimeras has then led to the identification and functional characterization of a new motif, in the CTD of HIV-1 group M integrase, essential for viral integration.

## Results

### Analysis of intergroup M/O chimeras in the CTD of IN

The functionality of the integrases studied in this work was evaluated following the protocol outlined in Figure 1A-B and detailed in Materials and Methods. For this, we replaced the original RT and IN sequences of the p8.91-MB (see Materials and Methods) by those of either one isolate of HIV-1 group M, subtype A2, referred herein as “isolate A” or one isolate of HIV-1 group O, referred herein as “isolate O”. The resulting vectors were named RTA-INA (vRTA-INA) and (vRTO-INO), respectively. With these vectors, we estimated the functionality of the integrases by measuring the efficiency of generation of proviral DNAs. Since the number of proviral DNAs generated for each sample is dependent not only on the levels of functionality of the integrase but also on the amount of total viral DNA generated after reverse transcription, we estimated the amount of total viral DNA generated by each sample by qPCR as described in Materials and Methods. In parallel, we measured the amount of proviral DNA generated either by the puromycin assay or by the Alu qPCR assay as described in Materials and Methods (Evaluation of integration by puromycin assay). The amount of proviral DNA divided by that of total viral DNA provides an estimate of the efficiency of integration. Comparable efficiencies of integration were measured with the two vectors, irrespective of whether the estimation was done using the puromycin assay (71 ± 13 % and 72 ± 24 % the level of the reference vector v8.91-MB respectively) or the Alu qPCR assay (70 ± 14 % and 68 ± 15%, respectively, Figure 1C). Throughout this work, the efficiency of integration has always been evaluated by the puromycin-resistance assay normalised by the amount of total DNA. Control vectors, in which the catalytic activity either of the integrase or of the reverse transcriptase have been abolished, in vRTA-INA, by the introduction of the D116A mutation in IN or of the D110N-D185N mutations in RT (43, 44), gave the expected results (Figure 1C).

**Figure 1.**
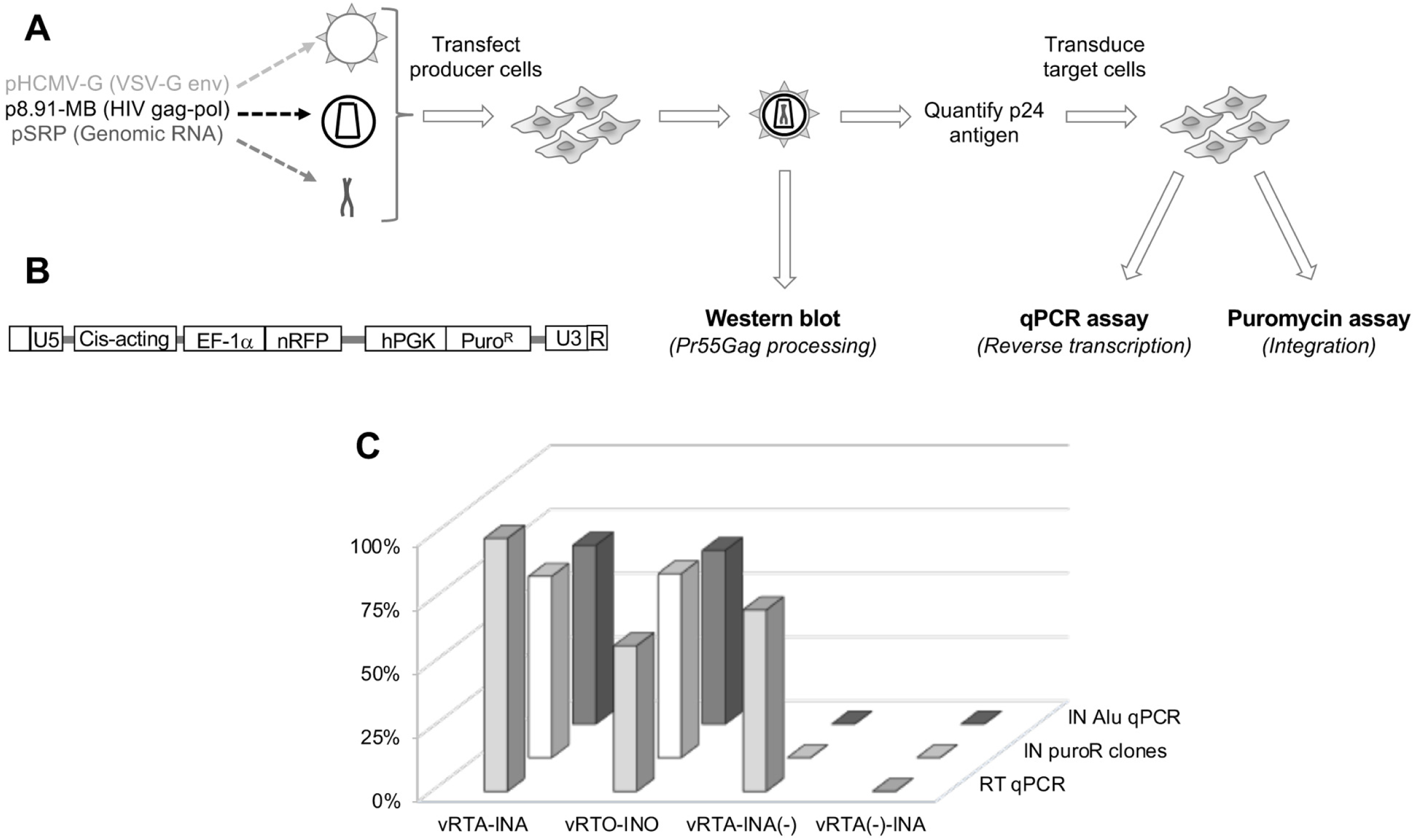
Outline of the experimental system. *Panel A.* Workflow used to evaluate Pr55Gag processing, reverse transcription, and integration in our experimental system. VSV-pseudotyped HIV-1 derived vectors, produced by triple transfection, were used to transduce HEK 293T cells. Upon integration, the proviral DNA will allow growth of the cellular clones in the presence of puromycin. For multiplicities of infection lower than 1, the number of clones obtained is directly proportional to the number of integration events. *Panel B.* schematic representation of the viral genomic RNA contained in the viral vectors, transcribed from pSRP (panel A, also see Materials and Methods). R, U5 and U3, viral sequences constituting the LTR; “cis-acting”, viral sequences required for RNA packaging and reverse transcription; EF1-α and hPGK, internal human promoters driving the expression of the nuclear RFP (nRFP) and of the puromycin N-acetyl-transferase that confers resistance to puromycin (Puro^R^), respectively. *Panel C.* Evaluation of reverse transcription and integration in control samples, compared to v8.91-MB reference vector. The results give the average values of three independent experiments.

We chose to probe the existence of functional motifs in the C-terminal domain of integrase because this domain is involved in several non-catalytic functions of the protein. The C-terminal domain of INO used in this study is 10 amino acids longer (212-298) than that of INA (212-288, Figure 2A). We constructed three chimeras between isolates A and O, named after the position, in amino acids from the beginning of the IN-coding region, where the sequence shifts from that of one isolate to that of the other (Figure 2B). Chimera A(1-212)-O(213-298) is constituted by INA with the entire CTD from INO; chimera A(1-285)-O(286-298) is INA with the additional 10 amino acids of INO at the C-ter end plus the two most C-ter different amino acids; finally, as the region between position 212 and 288 differs in 12 amino acids, chimera A(1-272)-O(273-298) was constructed in such a way as to split the 12 different amino acids in two groups of 6.

**Figure 2:**
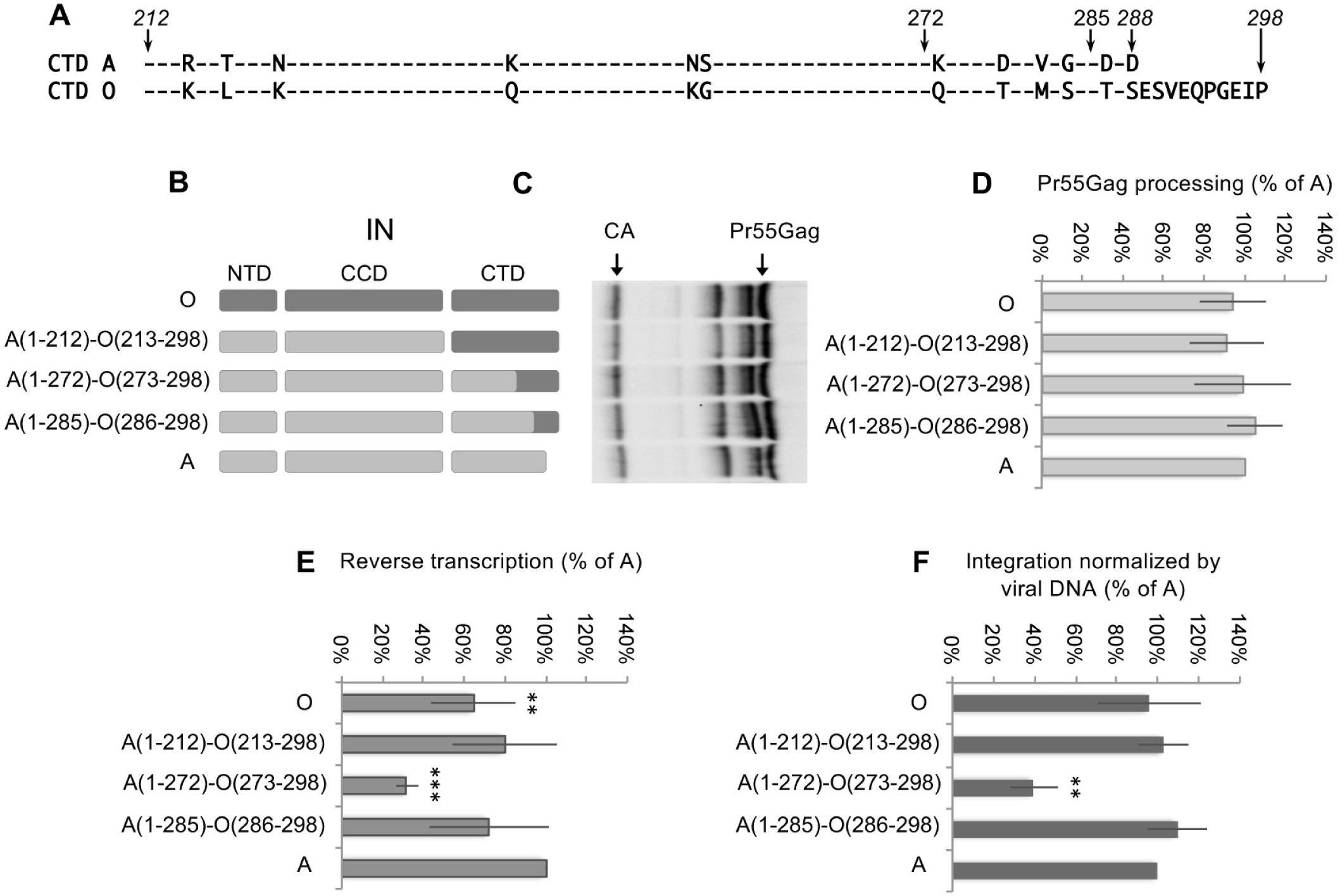
Functionality of chimerical integrases. *Panel A.* Alignment of CTD sequences from isolates A and O, used in this study. The numbers in italic on the left and on the right of the alignment indicate the beginning and the end (in amino acid) of the CTD, respectively. Only amino acids divergent between the two sequences are indicated by letters. The arrows and numbers above the alignment indicate the last position that, in the chimeras, was concordant with the sequence of isolate A. *Panel B.* Schematic representation of the integrases studied. Integrase from isolate O is drawn at the top of the panel in dark grey; integrase from isolate A is drawn at the bottom of the panel in light grey. The genetic origin of the portions of the chimeras is indicated by the colour code that refers to the reference isolates A and O. *Panel C.* Representative western blot obtained with an anti-CA mouse monoclonal antibody. *Panel D.* Efficiency of processing of the Pr55Gag precursor, estimated by the amount of CA compared to the amount of Pr55Gag precursors detected by western blot (as in panel B). The results are expressed as function of the reference wt IN A, set at 100%. *Panel E.* Efficiency of reverse transcription (detection of the junction U5-Psi by qPCR) expressed as function of the reference wt IN A. *Panel F.* Efficiency of integration calculated with the puromycin assay, normalized by the amount of total viral DNA (estimated by qPCR), expressed as function of the reference wt IN A. Error bars indicate standard deviations. The results given in panels C-E are the average of 3 independent experiments. ** p <0.01; *** p <0.001, p values for comparison to wt IN A.

We first performed western blots (Figure 2C) on viral particles to monitor the degree of proteolytic processing of the Gag precursor (Pr55Gag), since incomplete processing would result in immature viral particles, affecting infectivity. No significant differences in Pr55Gag were observed between isolate O and chimerical constructs compared to isolate A (Figure 2D). We then evaluated the efficiency of reverse transcription (measuring the amount of viral DNA produced by qPCR) and of integration (as described above). Only chimera A(1-272)-O(273-298) exhibited significant defects in both reverse transcription and integration (Figure 2E-F), suggesting that a covariation network, present between positions 212 and 285, was broken in this chimera. Since in these experiments the IN is expressed from p8.91-MB and not from the genomic RNA, it can be ruled out that the phenotypes observed are due to an effect of the mutations on the genomic RNA, as for example on the process of splicing, as it has been previously described for some mutants of the C-terminal domain of IN (45).

### Characterization of IN CTD

In order to evaluate the individual contribution of the 10 amino acids differing between positions 212 and 285 (Figure 2A), each residue in IN A was individually replaced by those of IN O and the ten-point mutants were tested for processing of Pr55Gag, reverse transcription and generation of integrated proviruses.

Except for mutant N254K, no significant difference in level of Pr55Gag proteolytic processing was observed between mutants and parental vector A (Figure 3A). The effect on reverse transcription was an overall reduction of efficiency for most of the mutants, with a residual efficiency between 45 and 90% that of the parental vector A (Figure 3B). Concerning integration efficiency, instead, the majority of the mutants did not show a significant decrease, except for mutants K240Q and K273Q for which integration was dramatically impaired (Figure 3C). This suggests a specific implication of these two residues in the integration process. When the two mutations were combined (K240Q/K273Q mutant), while the level of reverse transcription remained above 40 % that of wt IN A, integration dropped to undetectable levels (Figure 3D).

**Figure 3.**
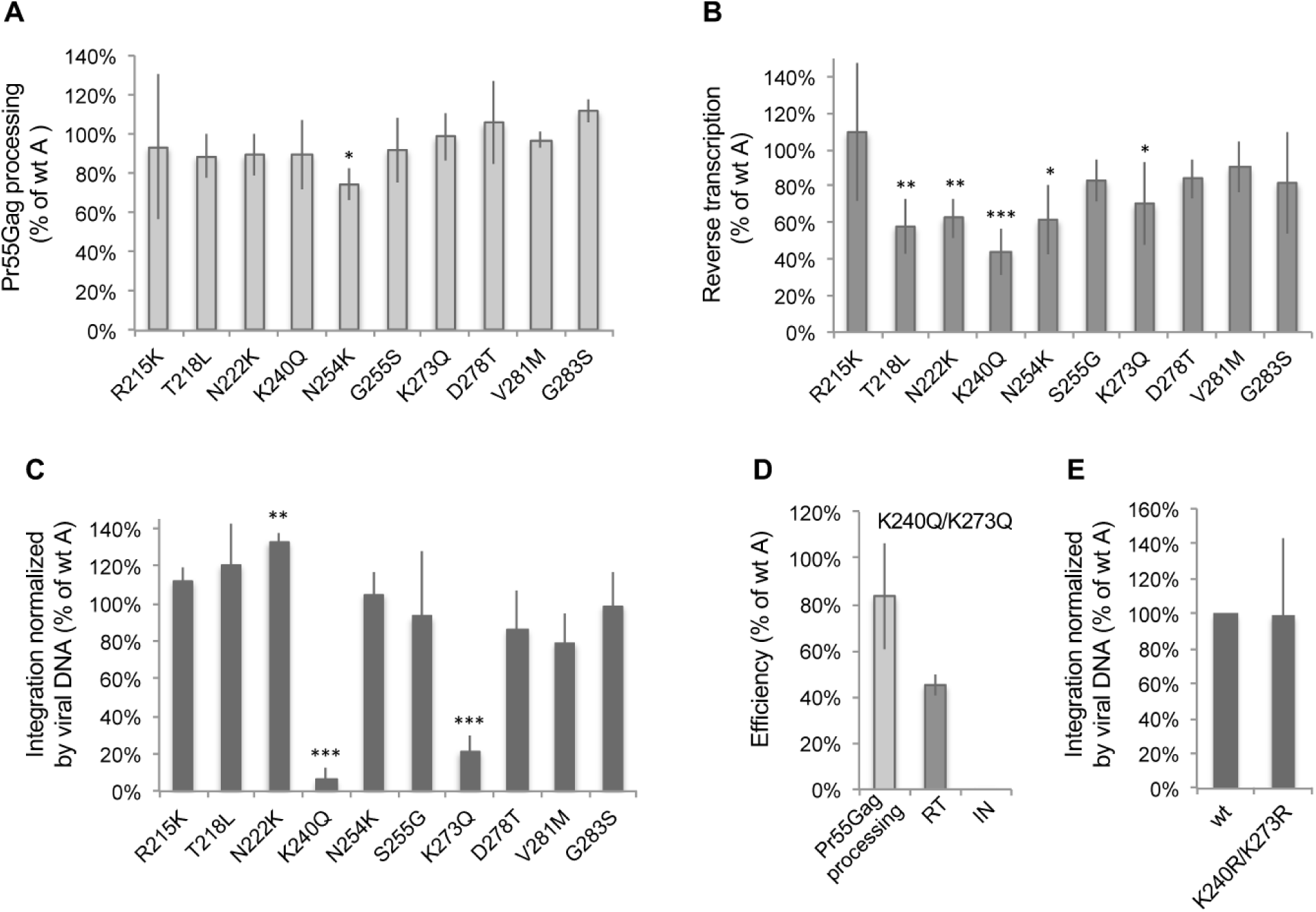
Functionality of IN A with mutated CTD. *Panels A-C*. Efficiency of processing of the Pr55Gag precursor (panel A), of reverse transcription (panel B), and normalized efficiency of integration (panel C). *Panel D*. Efficiency of processing of the Pr55Gag precursor, of reverse transcription, and of normalized efficiency of integration for the K240Q/K273Q mutant. *Panel E*. Efficiency of integration of the K240R/K273R mutant (NRNR in the Figure) and of wt IN A (*NKNK, set at 100%). Error bars indicate standard deviations. In all panels the results are the average of 3 independent experiments. * p <0.05; ** p <0.01; *** p <0.001, p values for comparison to wt IN A.

To discriminate between the role of the charge of K_240_ and K_273_ from that of their possible acetylation, we replaced both residues by two R (K240R/K273R mutant). The level of integration of this mutant was comparable to that of wt IN A (Figure 3E), indicating that the presence of a positive charge and not acetylation at these positions was important for integration efficiency. However, these mutations reduced by half both Pr55Gag proteolytic processing and reverse transcription (Figure S1A).

In both mutants showing a marked defect in integration (K240Q and K273Q), a K (positively charged polar side chain) was replaced by a Q (non-charged polar side chain), the amino acid present in isolate O at the corresponding positions. Conversely, in isolate O, two K are present in positions where a polar non charged amino acid (N in both cases) is present in isolate A (positions 222 and 254, Figure 2A). Therefore, in order to evaluate if also the two non-charged polar amino acids (N) present in isolate A at positions 222 and 254 are essential, we replaced them by a non-polar amino acid like leucine (mutant LKLK) and, in parallel, by a non-charged polar residue, Q (mutant QKQK). While in the LKLK mutant the efficiency of integration dropped to almost undetectable levels, in the QKQK one it was comparable to that of the wt enzyme, suggesting that the presence of a polar residue at these positions is essential (Figure 4A). To understand whether the polar nature of the amino acid at positions 222 and 254 is enough to retain functionality, the N where replaced by two threonine, which are polar but do not have the amide group of asparagine. In this case (TKTK mutant) integration dropped to undetectable levels (Figure 4A) indicating that not only the polarity is important but also the functional group carried by the amino acid. Therefore, the biochemical features of all four residues identified are important.

**Figure 4.**
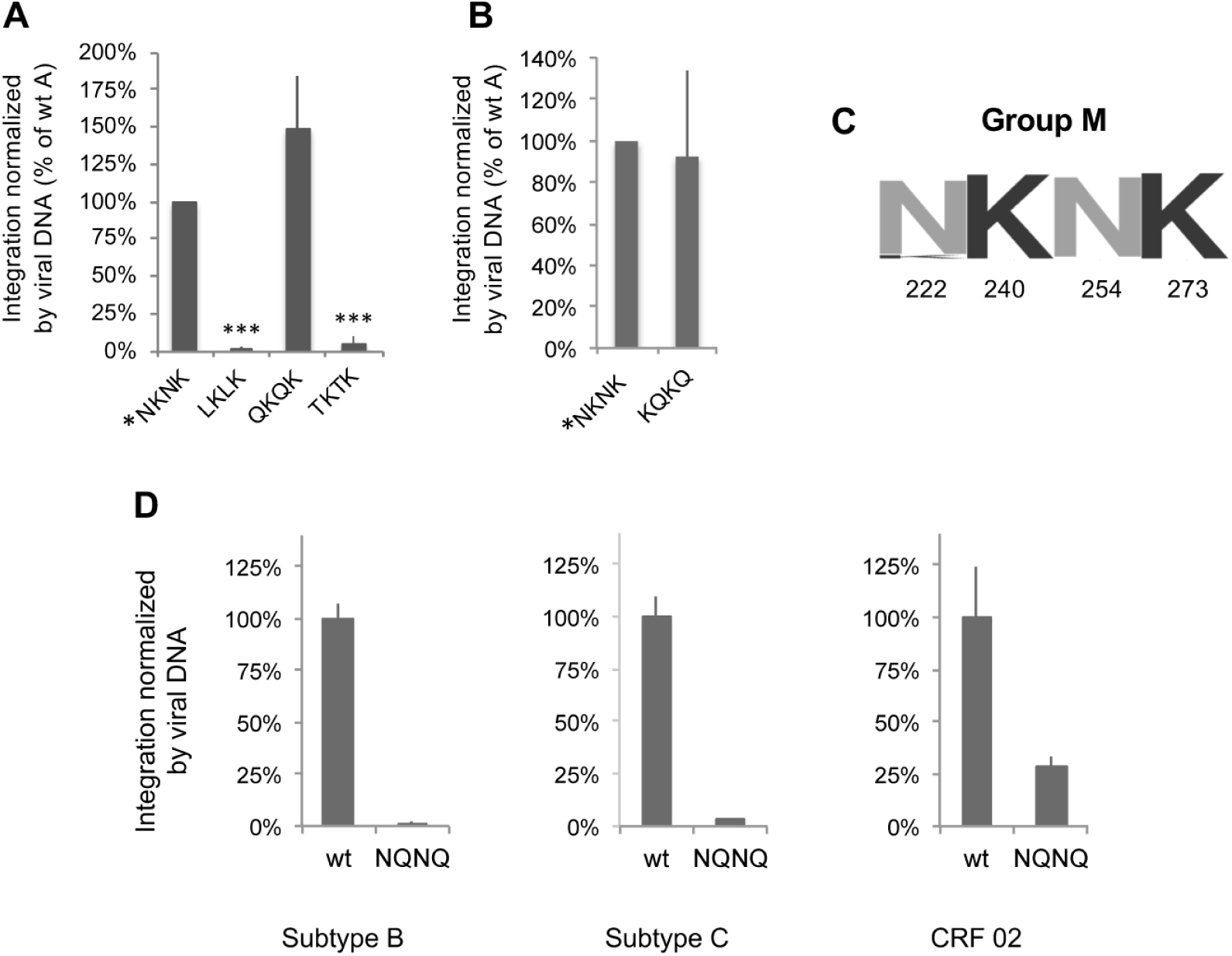
Definition of the NKNK motif and of its importance in the most widespread phylogenetic groups of HIV-1. *Panel A*. Efficiency of integration of various mutants of the N residues of the NKNK motif and of the wt enzyme (*NKNK, set at 100%). *Panel B.* Efficiency of integration of the mutant carrying the sequences of isolate O at positions 222, 240, 254 and 273 (K_222_Q_240_K_254_Q_273_, KQKQ in the Figure) and of the wt enzyme (*NKNK, set at 100%). *Panel C.* Conservation logo of the sequence at positions 222, 240, 254 and 273 in HIV-1 group M integrases. *Panel D.* Efficiency of integration of the double mutant N240Q/K273Q (NQNQ in the Figure) of an isolate of subtype B, one of subtype C and from CRF02, compared to the corresponding wt integrases, set as reference at 100 %. In all vectors the RT sequence had the same phylogenetic origin as IN and was replaced using the *Mlu*I-*Bsp*EI cassette in p8.91MB, as described in Materials and Methods. Error bars indicate standard deviations (standard deviations of the wt integrases of each subtype were calculated with respect to reference wt IN A, used as control). The results are the average of 3 independent experiments.

Finally, we wondered whether the residues present at positions 222, 240, 254 and 273 could be interchanged between isolates O and A. Therefore, we generated the quadruple mutant of isolate A N222K/K240Q/N254K/K273Q (called KQKQ for simplicity). Remarkably, the integration efficiency of this mutant was not significantly different from that of wt IN A (Figure 4B), indicating the existence of a functional link between these four positions.

The alignment of HIV-1 IN sequences reveals a strong conservation of the amino acids N_222_K_240_N_254_K_273_ in group M (Figure 4C). To confirm the need for K_240_ and K_273_, observed in isolate A, also for other isolates of group M, we introduced the K240Q/K273Q double mutation (NQNQ mutant) in integrases from three other primary isolates of group M (Figure 4D). In all cases, a dramatic drop in integration was observed with respect to the corresponding wt integrases, confirming the results obtained with isolate A. The importance of the two K in the motif was therefore confirmed in isolates from the most widespread HIV-1 group M subtypes in the epidemics, subtypes A, B, C, and CRF02 being responsible for 79% of the HIV-1 infections worldwide (46).

The possibility of permuting the positions of the four amino acids at positions 222, 240, 254 and 273 indicates a functional relationship between these residues that can therefore be considered as a functional motif that, based on the identity of the amino acids present at these positions in isolates of group M, we refer to as the “NKNK” motif.

### Importance of the lysines in the NKNK motif

To understand to which extent the number and the positions of the K in the motif influence IN functionality, we generated a series of mutants based on the replacement of the amino acids present in isolate A by those of isolate O. We thus tested all possible variants (Figure 5A) containing either only one K (four mutants, Figure 5B), two K (five mutants plus the wt, Figure 5C), three K (four mutants, Figure 5D) or four K (one mutant, KKKK, Figure 5A) at any of the positions in the motif. The presence of a single K led on average to a drop to 20% of integration with respect to wt IN A, whereas when two or more lysines were present in the motif, levels of integration were close to those of wt IN A, ranging from 75 to 137% (Figure 5A). The mutant with no K, where the motif sequence has been changed from NKNK to NQNQ, confirmed the total loss of integration already observed with this mutant (see Figure 3D). Finally, from the analyses of the different mutants it appears that the presence of a K at the first position of the motif (position 222) consistently leads to a higher level of integration in all classes of mutants (those with 1, 2, or 3 K). Interestingly, though, position 222 has a N in the wt enzyme.

**Figure 5.**
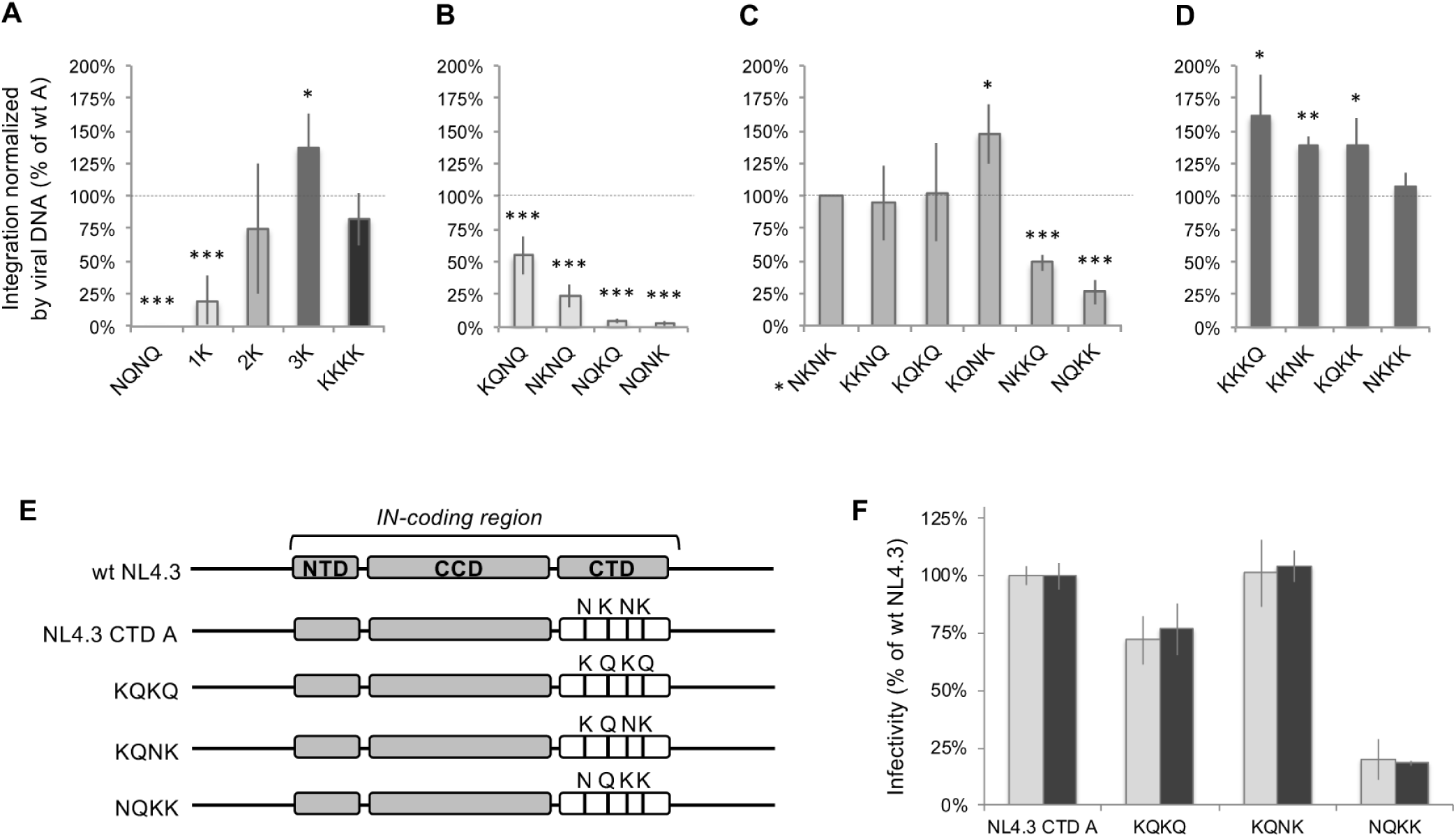
Importance of the number and position of the K residues in the N_222_K_240_N_254_K_273_ motif of the CTD. *Panel A.* Efficiency of integration, normalized by the amount of viral DNA, for the IN mutants grouped by the number of K present at positions 222, 240, 254, 273. The composition in amino acids in the four positions of the motif is given for the isolate with 0 and for the one with 4 K. For clarity, only the four letters of the amino acids of the motif are represented for each mutant, omitting the positions; the first letter indicates the residue at position 222, the second, position 240, the third, position 254 and the fourth, position 273. *Panels B-D*. Efficiency of integration of the individual mutants containing 1 (panel B), 2 (panel C), or 3 (panel D) K in the motif. In panel C, the motif corresponding to the sequence of wt IN A (reference set at 100%) is indicated by an asterisk. Error bars indicate standard deviations. The results are the average of 4 independent experiments. * p <0.05; ** p <0.01; *** p <0.001, p values for comparison to wt IN A. *Panels E-F*, importance of the NKNK motif in replication-competent viruses. *Panel E.* Scheme of the portion coding for the integrase in the various viruses. Drawn in grey are the parts derived from the NL4.3 sequences, in white those from isolate A. The black bars indicate positions 222, 240, 254 and 273 from left to right; the amino acid found for each mutant in each of these four positions is indicated above the bars. *Panel F.* Infectivity of the viruses shown in panel E (except wt NL4.3 that is used as reference, set at 100%). The results are given in grey for CEM-SS and in black for TZM-bL cells. Error bars indicate standard deviations with respect to the reference wt pNL4-3. The results are the average of 2 independent experiments.

When considering individual mutants within the different classes, we observed a significant decrease in functionality for all the mutants possessing only one K (Figure 5B). For the mutants containing two K, three variants were at least as functional as wt IN A (NKNK in the figure), while two displayed a significant reduction (Figure 5C). Finally, all mutants containing three K were at least as functional as the parental IN A (Figure 5D). Remarkably, the results obtained with the mutants containing three or four K indicate that the positively charged residues can replace the polar ones, while the reverse is not the case, as shown by the mutants with none or only one K. Overall, these results indicate that at least two K are required to have wt levels of integration, even if not all the positions in the motif are equivalent. Instead, all mutants impacted reverse transcription with a reduction to 40-80% of the wt IN A (Figure S2).

### The NKNK motif in replication-competent viruses

To confirm the observations made in the single infection cycle system, some mutants were then tested in a replication-competent system using NL4.3 as primary virus. Mutants of the class containing two K in the motif (the number of K found in circulating viruses) and with a marked phenotype were chosen for this analysis. Besides the wt A sequence, we chose three mutants that either retained integration (KQKQ and KQNK) or exhibited reduced integration (NQKK) (Figure 5C). To construct the four variants, we replaced the sequence of NL4.3 CTD by that of isolate A, either wt or carrying the KQKQ, KQNK or NQKK motifs (Figure 5E).

The infectivity of the virus carrying the whole CTD of INA instead of that of NL4.3 (called NL4.3 CTD A, Figure 5E) was comparable to that of wt NL4.3 virus, set as reference, indicating that the replacement of the whole CTD from NL4.3 by that of isolate A did not impact viral infectivity (Figure 5F). Regarding the mutants, the results well recaptured the observations made with a single infection cycle (Figure 5C): the infectivity was maintained for KQKQ and KQNK mutants while it was markedly decreased with NQKK motif (Figure 5F).

### Role of the lysines of the motif in the integration process

In order to characterise in which steps of the infectious cycle are involved the lysines of the NKNK motif, we evaluated the effect of their mutation in two steps (other than reverse transcription) upstream the integration of the pre-proviral DNA in the chromosomes of the host cell. In particular, by quantifying the two LTR circles (2LTRc), we evaluated nuclear import and, by characterizing the LTR-LTR junctions of 2LTRc, the efficiency of 3’ processing, which takes place in the cytoplasm, before nuclear import.

2LTRc are exclusively formed in the nucleus and are, therefore, useful markers for nuclear import of the reverse transcribed genomes (47). They are generated when the full-length reverse transcription products are not used as substrate for integration. If a mutant is defective in catalysis but carries out nuclear import efficiently (as mutant D116A), 2LTRc should accumulate with respect to a wt IN. Instead, if the mutant is also impaired in nuclear import, 2LTRc will either not increase with respect to the wt IN or increase but more modestly than for D116A.

Hence, to monitor nuclear import, we measured the amount of (2LTRc) in wt IN A, in mutants containing either no K (NQNQ) or only one (either K_273_, NQNK mutant, or K_240_, NKNQ mutant). IN A D116A mutant was used as a control. This mutant being totally inactive for integration was considered to produce the highest accumulation of 2LTRc, set at 100%. As expected, the level of 2LTRc found with wt IN A, which efficiently imports and integrates the reverse transcribed genome, was significantly lower (25%) than that of the D116A mutant. As shown in Figure 6A, despite their inability to generate proviral DNA, the mutants had levels of 2LTRc significantly lower than IN A D116A, indicative of a defect in nuclear import.

**Figure 6.**
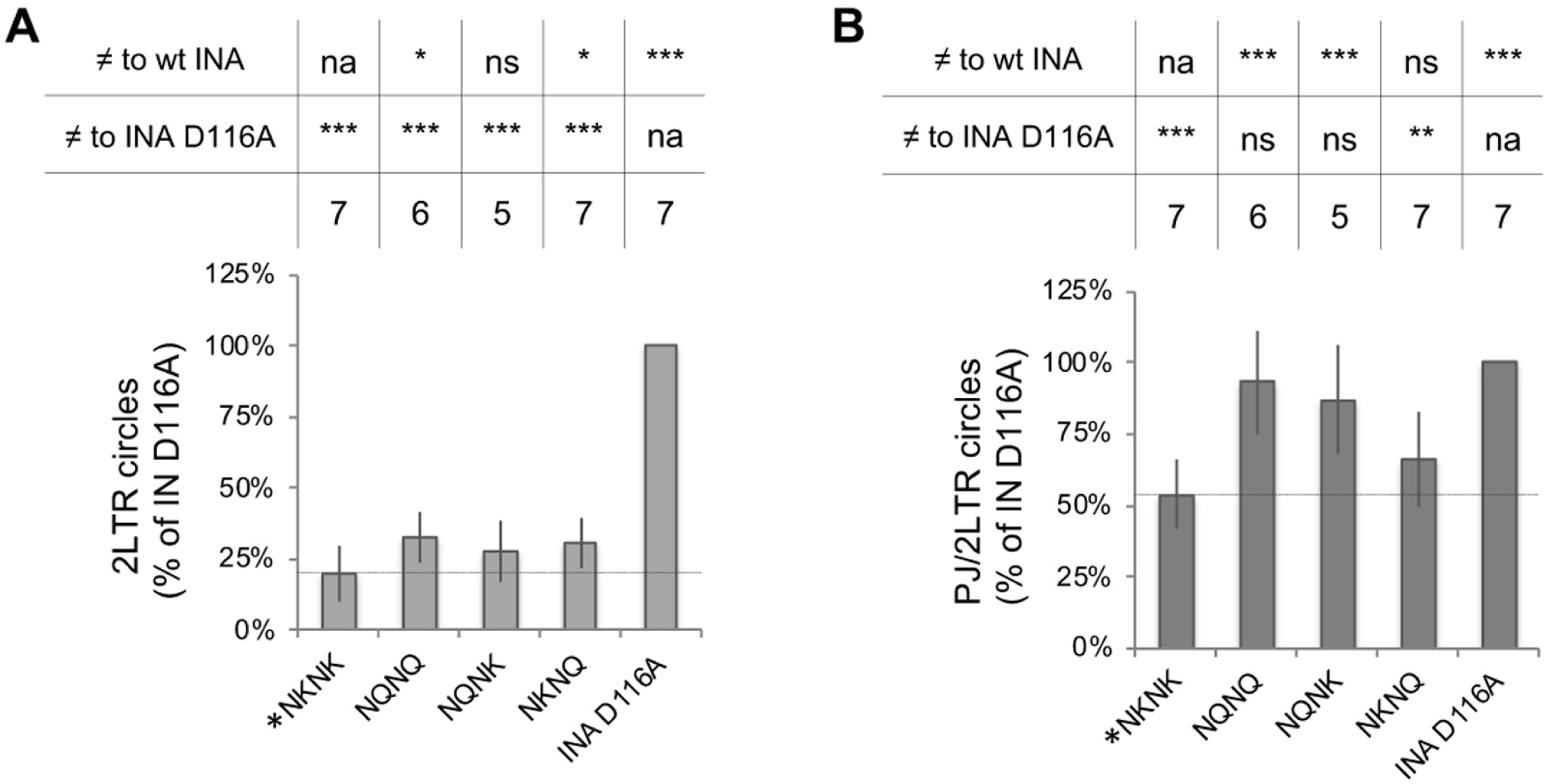
Amount of 2LTRc (*panel A*) and of ratio of PJ/2LTRc (*panel B*) in the mutants deprived of one or both K of the NKNK motif. The motif corresponding to the sequence of wt IN A (reference set at 100%) is indicated by an asterisk. Error bars indicate standard deviations. Above the plot are given the p values for the comparisons of the different samples with respect to wt IN A or to the integration-deficient mutant IN A D116A (* p <0.05; ** p <0.01; *** p <0.001). The number of independent experiments performed for each sample (n) is also given.

To estimate the efficiency of nuclear import in the mutants, we estimated the level of 2LTRc (Table 1, line 2, “theoretical level”), that could be obtained if no defect in nuclear import was present. We then calculated the efficiency of nuclear import as the ratio between the level of 2LTRc observed experimentally (Table 1, line 3) and the theoretical one. If no defects in nuclear import are present, ratios should be around 1, while defects in nuclear import would yield ratios <1. The ratios found for the three mutants were in the 0.31-0.35 range (Table 1, line 4), indicative of a reduction of nuclear import to approximately 1/3 that of the wt enzyme. Therefore, the defects in nuclear import contribute to the decrease in integration found with these mutants, but cannot alone account for the low levels observed, particularly in NQNQ and NQNK mutants for which integration was almost undetectable (Table 1, line 1).

**Table 1.**
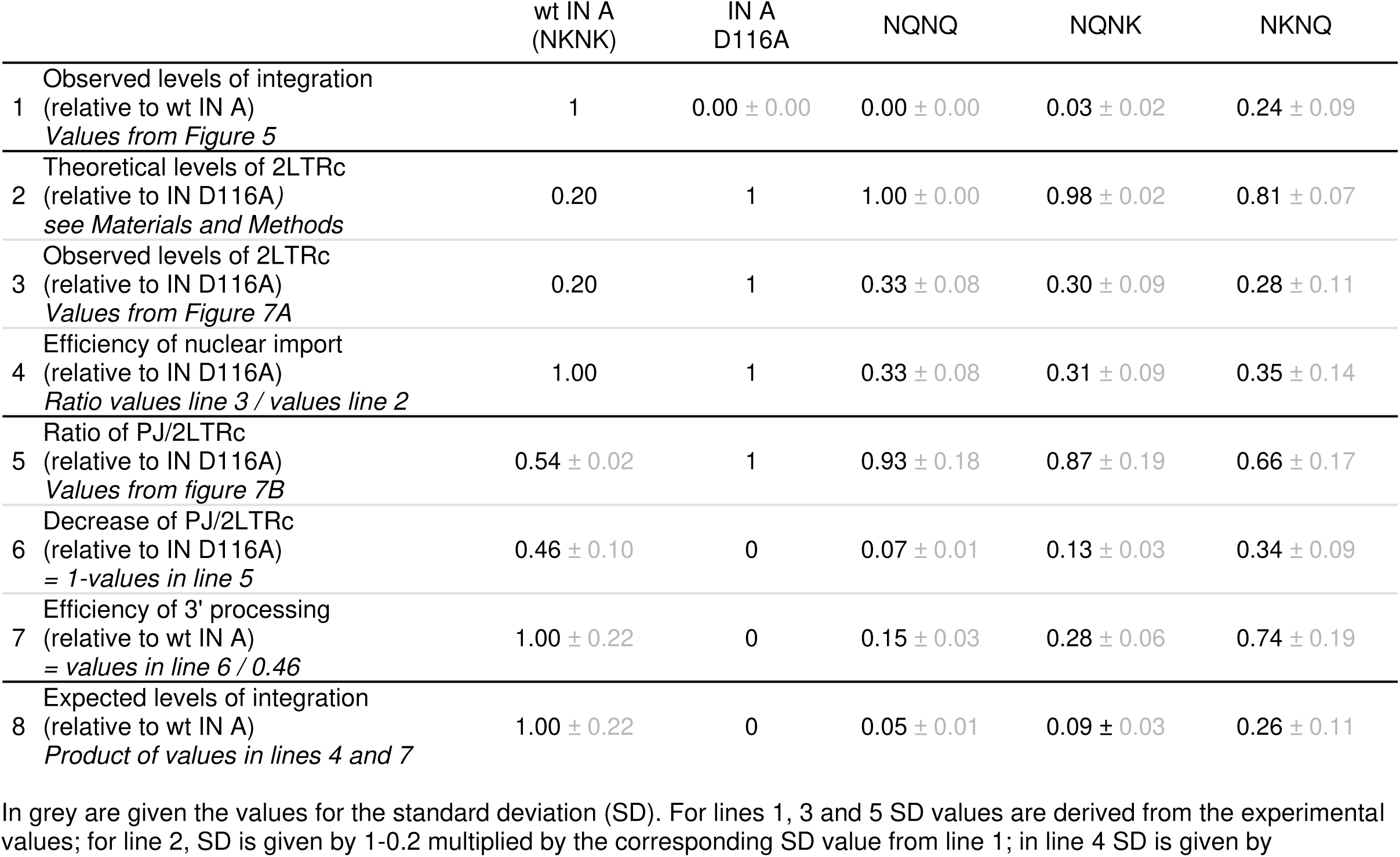
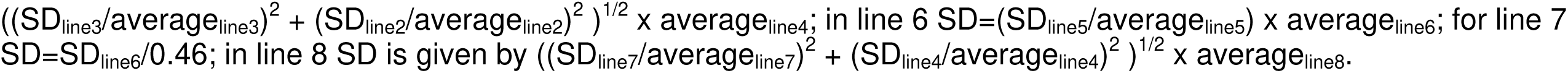
Estimate of the contribution of the defects of nuclear import and 3’ processing to the efficiency of integration observed with the mutants of the NKNK motif.

The efficiency of 3’ processing carried out by IN was then analysed by quantifying the different LTR-LTR junctions in the 2LTRc. The 2LTRc found in the nucleus are generated from DNAs carrying either unprocessed or processed 3’ ends. In the first case, the 2LTRc will present “perfect junctions” (PJ) while in the second the junctions will be “imperfect”. A high ratio of PJ/2LTRc is therefore indicative of inefficient 3’ processing. We found that mutating both K (mutant NQNQ) or only K_240_ (mutant NQNK) led to results not significantly different from those obtained with the IN A D116A catalytic mutant (Figure 6B), indicative of a marked defect in 3’ processing. Mutating K_273_ (mutant NKNQ), instead, did not affect the process, with PJ/2LTRc values comparable to those of the wt enzyme.

To evaluate the contribution of defects in 3’ processing to the decreased integration efficiency observed with the various mutants, we first estimated the maximum diminution of PJ/2LTRc ratio observed for a fully competent enzyme (wt IN A). The PJ/2LTRc ratio for wt IN A was 0.54 that of IN A D116A (Table 1, line 5), corresponding to a reduction of 46 % due to 3’ processing (Table 1, line 6). The ratio PJ/2LTRc with respect to IN A D116A was then calculated for each mutant and the resulting value was divided by 0.46, obtaining an estimate of the efficiency of 3’ processing relative to that observed for wt IN A (Table 1, line 7). 3’ processing of NQNQ and NQNK mutants was dramatically reduced, 15 % and 28 % that of wt IN A, respectively. Mutating K_273_, instead, only decreased 3’ processing to 74 % that of wt IN A.

Finally, in order to understand if these two types of defects (nuclear import and 3’ processing) were sufficient to explain the integration defects observed with the various mutants, we combined the effect of these defects (Table 1, line 8). The values obtained account remarkably well for the efficiencies of integration observed (lines 1 and 8 of Table 1) indicating that the decrease observed when mutating the K of the NKNK motif, once normalized for the differences observed in the amount of viral DNA produced, is essentially due to alterations in these two processes.

### Structural analysis of wt and mutant integrase C-ter domains

To understand the structural bases for the functional differences observed in the NKNK motif mutants, the crystal structures of the C-terminal domain (IN CTD, 220-270) of wild type IN A and of the reference strain NL4.3 were solved at 2.2 Å and 1.3 Å of resolution, respectively. For both structures, K_273_ was not included as it is in a disordered region of IN. For all crystal forms we observed a strong packing interaction through the His-tag coordinating a Nickel ion (Figure S3A). The quality of the structures and maps is shown in Figure S3, panels B-D. The structures had the same topology, consisting in a five-stranded β-barrel (Figure S4A). The region encompassing the positions of the motif (Figure 7A-B) generates a surface endowed with a positive potential (circled in yellow in Figure 7C-D), suggesting that this feature could be important for the functionality of the IN. In this case, it is expected that inserting additional lysines in the motif (as for the mutants containing three or four lysines) would retain functionality and, conversely, removing the K (mutants with one K or no K) would affect it. This is what we observed in Figure 5. Nevertheless, the correlation between surface potential and functionality is less clear for the mutants where the number of K in the motif (two) is not altered but their positions are permutated with polar amino acids.

**Figure 7.**
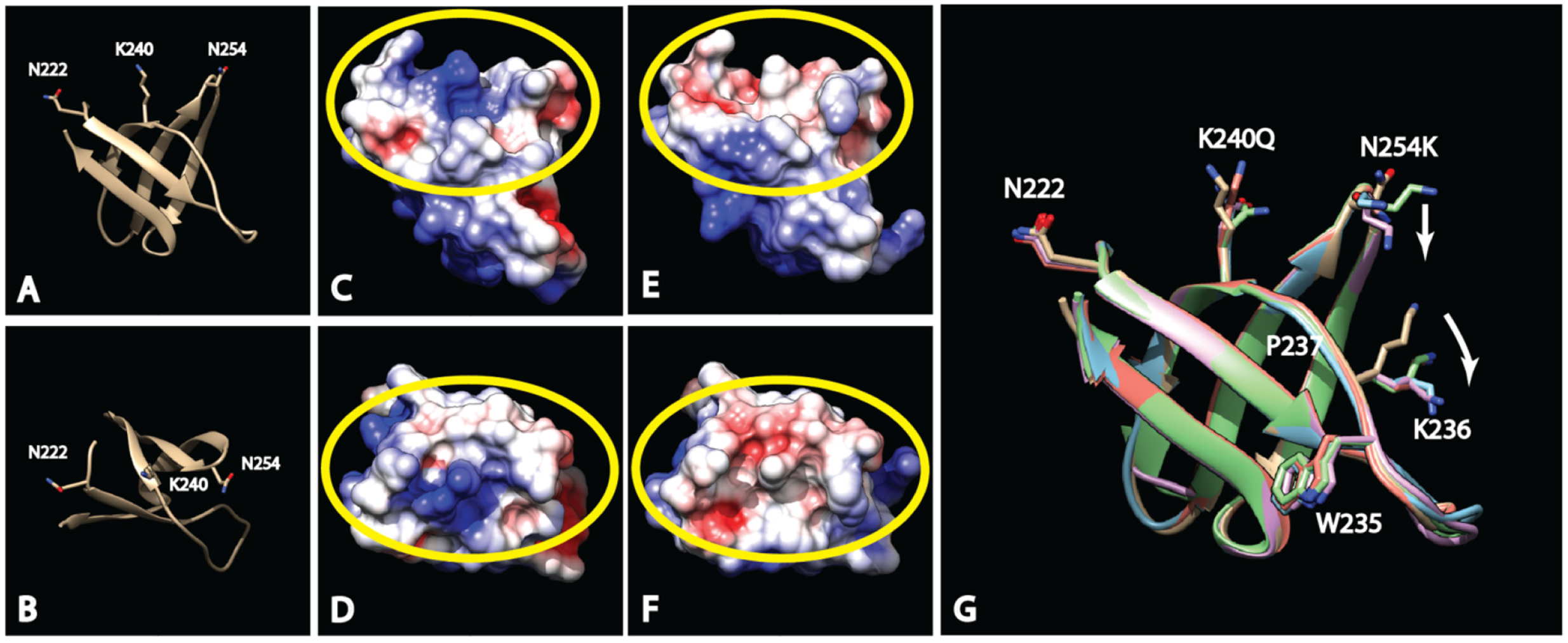
Structural analysis of the NKNK motif. *Panels A and B*. Side view (A) and top view (B) of the ribbon representation of the crystal structure of CTD A. The positions of residues N222, K240 and N254 are represented with sticks as well as the position of the I234, the only different residue between the CTD of IN A and IN NL4.3. *Panels C* (side view) *and D* (top view) are the surface electrostatic potential representation of CTD A. In red, negative potential; in blue, positive potential and in white neutral regions. Circled in yellow is the region with large differences in the mutant structures (see below *Panel H-M*). *Panel E*. Superposition of the CTDs of IN A and IN A NQKK (chain A, B and C). The mutation N254K induces a displacement of the K236 side chain (white arrows) disturbing the structure of the 235-237 region. *Panels F and G*. Side view (F) and top view (G) of the superposition of the three molecules in the asymmetric unit of the NQKK CTD. The position of the residue N222, K240Q mutation and N254K mutation are represented with sticks as well as the position of the I234. *Panels H, J and L* (side view) and *I, K and M* (top view) are the surface electrostatic potential representation of NQKK CTD chain A (H, I), chain B (J, K) and chain C (L, M). In red, negative potential; in blue, positive potential and in white neutral regions. The regions with large differences are circled in yellow.

To clarify this point, we solved, at a 2.0 Å resolution the crystal structure of the CTD of the NQKK mutant, which was the one displaying the most dramatic drop in integration among the mutants possessing two K (Figure 5C). The NQKK CTD crystalized in a different space group and had three chains in the asymmetric unit. The superposition of the five structures (Figure S4B) corresponding to NL4.3 CTD, to A CTD and to the three molecules in the asymmetric unit for the NQKK CTD (chains A, B and C) did not show significant differences in the main chain fold (Root-mean-square deviation of atomic positions, RMSD: NL4.3 CTD vs A CTD = 0.395 Å, A CTD vs NQKK CTD ABC = 0.653 Å, NQKK CTD chain A vs chain B vs chain C = 0.558 Å). Interestingly, the positive surface electrostatic potential observed for the wt enzyme was markedly perturbed in the NQKK mutant (yellow circle in Figure 7E-F), a change that could well account for the decrease of functionality of the NQKK mutant integrase.

To further analyse the impact of the mutations on the structure, we defined the regions which are naturally disordered (Intrinsically Disordered Regions, IDRs) by the superposition of the three molecules in the asymmetric unit of the NQKK CTD structure. We assume that the change in the RMSD obtained among the three molecules represents the natural IDRs. For the main chain, disordered regions with high RMSD are 228-232 and 243-248 (Figure S5A). These same regions are found to be similarly disordered when comparing the main chain RMSD of the C-terminal domain of IN A to that of IN A NQKK chains A, B and C (Figure S5B), indicating that the mutations have no effect on the C-alpha backbone fold of IN CTD.

The three structures were then analysed from the standpoint of the arrangement of the side chains. Calculating side chains RMSD, disordered portions were found to correspond to regions 222-225, 228-232 and 243-248 (Figure S5C). The comparison of A CTD and NL4.3 CTD, which differ between positions 220-270 only by a single amino acid change (V234 in NL4.3 replaced by I234 in A), expectedly, did not reveal significant differences between the two structures. Instead, when comparing A CTD to NQKK CTD A, B and C side chain deviation, we observed a difference in the structure of region 235-237 (Figure S5D). This difference appears to be due to the N254K mutation that induces a displacement of the side chain of lysine 236 (white arrows in Figure 7G). This is likely due to a repulsive interaction between the side chains of the two lysines, resulting in a perturbation of the structure in the 235-237 region.

To evaluate the importance of charge configurations in the context of the C-terminal domain of retroviral integrases, we performed an analysis of the electrostatic charge surface potential for other lentiviruses as well as for other retroviruses. Integrases C-terminal domain structures are available for HIV-1 A2, PDB 6T6I (this publication); HIV-1 PNL4.3, PDB 6T6E (this publication); SIV, PBD 1C6V (48); MVV, PDB 5LLJ (49); RSV, 1C0M (50); MMTV, PDB 5D7U (51); MMLV, PDB 2M9U (52); PFV, PDB 4E7I (53). First, we superposed the available structures. The superposition shows that they share a common fold (Figure 8A) as well as a low Root Mean Square Deviation on secondary structure backbone despite a very low sequence identity for some cases (Table S1). A structure-based sequence alignment has then been performed (Figure 8B). Surprisingly, despite a low overall sequence identity (10 – 20 %) for some integrases, several regions have a strong local sequence similarity (red and yellow background in Figure 8B) while no conservation is observed at positions 222, 240 and 254 among lentiviruses nor retroviruses. However, when we compared the electrostatic surface potentials of all structures (Figure 8, Panels C-L), we could define two general retroviral classes. A first class represented by lentiviruses where the surface corresponding to the one delimited by the NKNK motif in HIV-1 M is basic and a second class represented by the other retroviruses tested (orthoretroviruses α, β, γ, and spumaretroviruses) where this surface is acidic or neutral.

**Figure 8.**
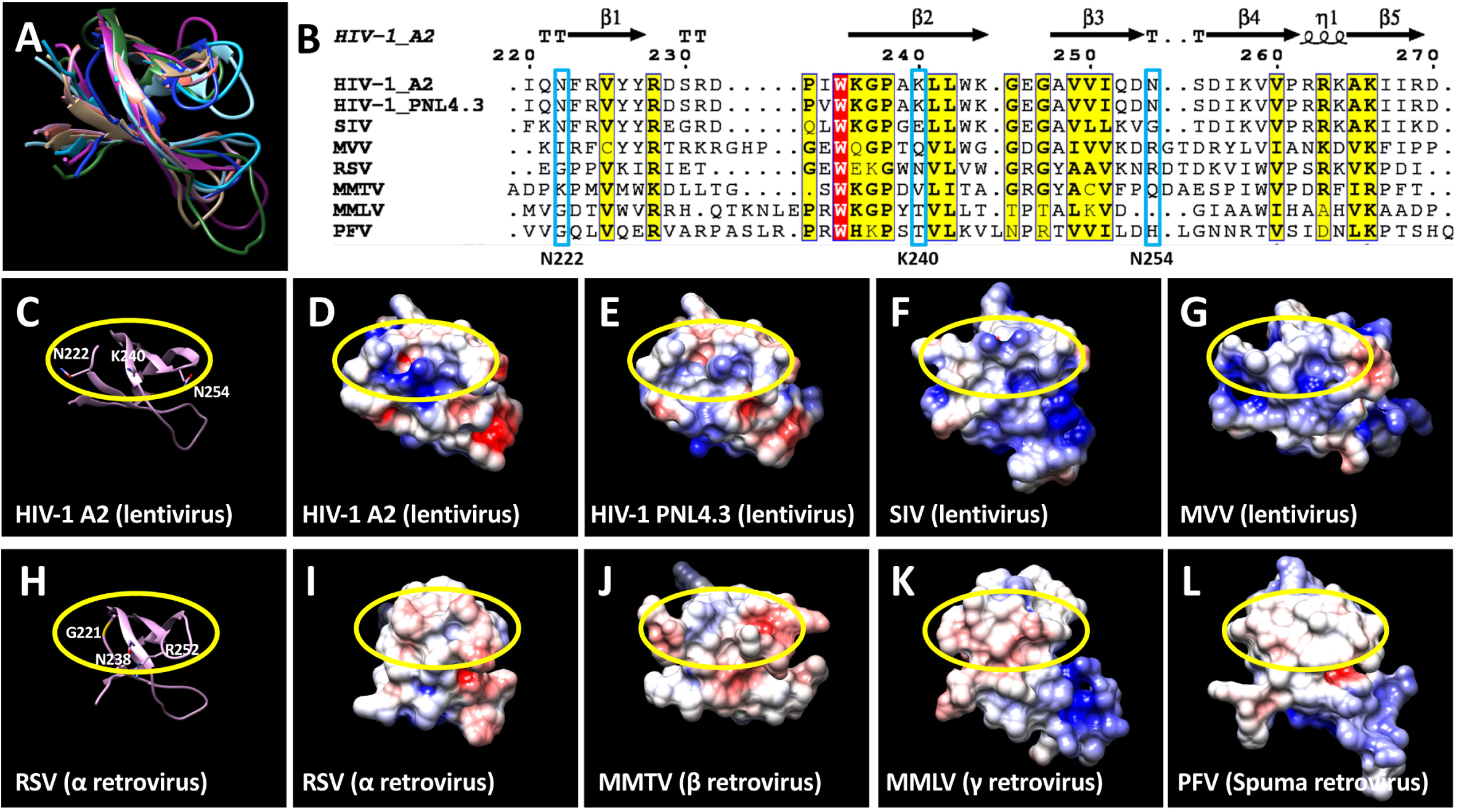
Analysis of the surface electrostatic potential in the C-ter of retroviral integrases. The structures and the sequences of the C-terminal domains have been extracted from: HIV-1 A2, PDB 6T6I (this publication); HIV-1 PNL4.3, PDB 6T6E (this publication); SIV, PBD 1C6V (48); MVV, PDB 5LLJ (49); RSV, PDB 1C0M (50); MMTV, PDB 5D7U (51); MMLV, PDB 2M9U (52); PFV, PDB 4E7I (53). *Panel A.* Superposition of the structure of the integrase C-terminal domains of four lentiviruses (HIV-1 A2, pink; HIV-1 pNL4.3, orange; Simian Immunodeficiency Virus, SIV, khaki; Maedi-Visna virus, MVV, cyan), of an α retrovirus (the Rous Sarcoma Virus, RSV, blue), of a β retrovirus (the Mouse Mammary Tumor Virus, MMTV, sky blue), of a γ retrovirus (the Moloney Murine Leukemia Virus, MMLV, purple), and of a spumaretrovirus (the Prototype Foamy Virus, PFV, green). *Panel B.* Structure-based sequences alignment of the integrases C-terminal domains. Sequence numbering corresponds to the HIV-1 A2 integrase sequence. Secondary structures from HIV-1 A2 are represented (TT: β-Turn, β1 to β5: β-sheets, η1: 3_10_-helix). Residues framed in blue: Position in the alignment of the three first amino acids from the NKNK motif. Red background: 100% identity in the sequence alignment. Yellow background: % of equivalent residues > 70% (considering their physical-chemical properties), equivalent residues are depicted in bold. *Panels C-L.* Surface electrostatic potential representation of integrases from several retroviruses. The surface corresponding to that delimitated by the NKNK motif in HIV-1 M (panels C, D and E) is circled in yellow. *Panel C*, ribbon representation of HIV-1 A2 C-terminal domain structure. The amino acids belonging to the NKNK motif are represented in sticks. *Panels D-G.* Surface potential representation of the C-terminal domain structures of four lentiviral integrases: HIV-1 A2 (panel D), HIV-1 pNL4.3 (panel E), Simian Immunodeficiency Virus (SIV) (panel F) and Maedi-Visna virus (MVV) (panel G). *Panel H.* Ribbon representation of the structure of Rous Sarcoma Virus (RSV) C-terminal domain. The surface circled in yellow corresponds to that delimitated by the NKNK motif in HIV-1 M after superposition of the structures. The amino acids corresponding to the motif in the structure-based alignment are shown as sticks. *Panel I.* Surface potential representation of the C-terminal domain structure of an α retrovirus, the Rous Sarcoma Virus (RSV). *Panel J.* Surface potential representation of the C-terminal domain structure of a β retrovirus, the Mouse Mammary Tumor Virus (MMTV). *Panel K.* Surface potential representation of the C-terminal domain structure of a γ retrovirus, the Moloney Murine Leukemia Virus (MMLV). *Panel L.* Surface potential representation of the C-terminal domain structure of a spumaretrovirus, the Prototype Foamy Virus (PFV). Negative potential is in red, neutral in white and positive potential is in blue.

## Discussion

Here, by performing a systematic comparison between the non-conserved amino acids in the CTD of the HIV-1 group M and group O integrases, we identify, in group M, a highly conserved motif that is essential for integration. The motif is constituted of two asparagines and two lysines (N_222_K_240_N_254_K_273_) all required for the generation of proviral DNA. In particular, when the K were mutated, integration was abolished due to the cumulative effects of decreased 3’ processing and nuclear import of the reverse transcription products (Table 1). Replacing the K by R did not affect integration (Figure 3E), suggesting that the essential feature of the K is their positive charge. Importantly, the positions of the two K of the motif could be permutated without affecting the functionality of the integrase in most cases (Figure 5).

A potential explanation for the retention of functionality when permutating the positions of the K across the motif comes from the structural data on the CTD obtained in this work. We have solved the crystal structure of the CTD of the wt IN A used in this study as well as that of the K240Q-N254K mutant (referred as NQKK in the result section). In the structure of the wt enzyme, the residues constituting the motif (except K_273_ which is part of an unresolved region) generate a positively charged surface (Figure 7A-D). This positive electrostatic potential surface is absent, instead, in the NQKK mutant (Figure 7E-F) which, despite the presence of two K, displays a drastic reduction of integration efficiency. These results, combined to the tests of functionality of the different mutants, suggest that the relevant parameter is the presence of a positive charge across this surface. Charged residues have a strong effect on the surface potential. The nature of the amino acid side chain (charged, polar, non-polar) on the surface of the protein defines the surface potential. Charged and polar groups, through forming ion pairs, hydrogen bonds, and other electrostatic interactions, impart important properties to proteins. Modulation of the charges on the amino acids, e.g. by pH and by post-translational modifications, have significant effects on protein – protein and protein – nucleic acid interactions (54). In addition to residues carrying net charges, also polar residues have significant partial charges and can form hydrogen bonds and other specific electrostatic interactions among themselves and with charged residues (55). In the case of the present study, the possible contribution of these mechanisms to the functionality of the integrase could be reflected by the loss of functionality observed by replacing the N, which carries an amide side chain (–CONH_2_), by either a non-polar amino acid (L) or by a polar one (T) but carrying a hydroxyl side chain (–OH). The analysis of the electrostatic surface potential for the integrases C-terminal domain of the retroviruses for which this is known showed that, despite a low sequence identity among some of the retroviruses (Table S1), the topology of the structure is maintained (Figure 8A) and the analysis of the surface electrostatic potential splits the viruses studied in two classes. One, constituted by lentiviruses, for which the surface delimited by the NKNK motif of HIV-1 M contains basic charges (in some cases brought by amino acids non corresponding to those of the NKNK motif of HIV-1 M). The second class including the other orthoretroviruses studied (α, β, γ) and spumaretroviruses where this surface contains acidic and neutral regions. This presence of basic regions, specifically in lentiviruses, could contribute to some specificity of lentiviral biology as to increase the efficiency of infection of quiescent cells.

The importance for protein functionality of charge configurations and clusters in their three-dimensional structures has been underlined by several studies (56-59). Charge permutations have been used in the NC region of the Gag protein for the Mason-Pfizer Monkey Virus (60). This basic region could be replaced with nonspecific sequences containing basic amino acid residues, without altering its functionality while mutants with neutral or negatively charged residues showed a large drop in viral infectivity in single round experiments. Moreover, a mutant exhibiting an increased net charge of the basic region, was 30% more infectious than the wild type. Also, in our study, increasing the positive charge of the NKNK motif of HIV-1 IN by introducing a third K leads to a slight increase in integration with respect to wt IN (Figure 5A and D).

As retention of IN functionality relies on the electrostatic surface potential rather than on the specific positions of the positively charged amino acid, we infer that this region is probably involved in the interaction with a partner carrying a repetitive negatively charged biochemical motif, as the phosphates of the nucleic acids backbone. Alternatively, the partner could be a disordered region of a protein that can rearrange to preserve the interaction when the positions of the positive charges are permutated across the surface of the NKNK motif. Indeed, the molecular recognition between charged surfaces and flexible macromolecules like DNA, RNA and intrinsically disordered protein regions has been observed for the Prototype Foamy Virus and Rous Sarcoma Virus Gag precursors (61, 62), for EBNA proteins of the Epstein-Barr virus (63), UL34 protein of the Herpes Simplex Virus (64) as well as for cellular proteins like APOBEC3G (65). Moreover, the presence of asparagines in the motif, which we show are required for integrase functionality, could contribute to the interaction with the nucleic acid or with a protein partner through hydrogen bonds with the bases (54) or with polar amino acids (55, 66, 67), respectively. The analysis of the motif in the context of the well-characterised structure of the intasome (31), mimicking a post-integration desoxyribonucleic complex, indicates that the residues forming the electrostatic surface point toward the solvent, at the exterior of the structure (Figure S6). This is coherent with the observation that mutating the motif does not affect late steps of the integration process, but rather earlier ones as 3’ processing and nuclear import.

We show that the NKNK motif of the CTD is involved in 3’ processing and nuclear import of the reverse transcription product. Indeed, the removal of the K impacts both processes and when combined, these effects are sufficient to account for the drop of infectivity to the undetectable levels observed in the absence of K in the motif (Table 1). Since mutating K_240_ has a strong impact on both 3’ processing and nuclear import, while K_273_ appears to be predominantly involved in nuclear import, it is possible that the involvement of the motif in these two processes implicates structurally distinct functional complexes. This is the first finding of an implication of the HIV-1 IN CTD in 3’ processing. So far, only the involvement of the catalytic domain had been demonstrated (39, 68, 69). Since it has been shown that different oligomerization states of IN influence specifically the ability to carry out 3’ processing or strand transfer (70), it is possible that the electrostatic surface formed by the NKNK motif help stabilize the oligomeric state that allows 3’ processing.

Concerning nuclear import, it is known that HIV-1 IN binds several cellular nuclear import factors through basic amino acids of the CTD, and that abolishing these interactions leads to non-infectious viruses displaying a severe defect in nuclear import. Here, we extend the regions of IN involved in this process by describing the need for a new motif, although it cannot be discriminated whether its involvement is direct or mediated by the interaction with a partner protein with karyophilic properties.

Some of the residues constituting the N_222_K_240_N_254_K_273_ motif have been previously characterized showing their implication in different steps of the infectious cycle. One is the involvement in reverse transcription. The integrase CTD interacts with the reverse transcriptase to improve DNA synthesis (71, 72). In one study, the double mutation K240A/K244E caused a decrease in reverse transcription to around 20 % the levels of the wt enzyme (72) while the K244E mutation alone caused a reduction of 40 % of RT efficiency (73), suggesting that K_240_ also contributes to reverse transcription. The decrease we observed when mutating K_240_ alone, to around 45 % of wt activity, is consistent with this view. The characterisation by NMR of the RT-binding surface in the IN CTD, obtained using the CTD_220-270_, shows that it is made up of 9 residues (amino acids 231-258 among which K_244_) that strongly interact both with the RT alone (9) and with the RT/DNA complex (74). When the interaction involves the complex, this surface includes 5 additional amino acids (74). Among these additional residues are N_222_ and K_240_, which are located at one edge of the surface. It is therefore possible that the nature of the residues at positions 222 and 240 affects the interaction between the CTD and RT/DNA complex.

Concerning K_273_, contradictory results have been obtained for reverse transcription of viruses harbouring integrases with sequential C-ter deletions (IN_1-270_ and IN_1-272_) (75, 76). Furthermore, for reverse transcription to occur, the genomic RNA must be encapsidated in the core of the viral particle. In this sense, it has been recently shown that mutating K_273_ together with R_269_ (R269A/K273A mutant) impairs encapsidation of the genomic RNA (6). As expected, reverse transcription in the double mutant was almost abolished. Here, mutating K_273_ to Q led only to a reduction of reverse transcription of 30% (Figure 3B), suggesting that mutating K_273_ alone is not sufficient to affect genomic incorporation into the viral capsid, at least in the majority of the particles. Supporting this, an earlier study (77) showed that the K273A single mutation did not affect viral replication in Jurkat cells, indicating that is the specific combination of R269A/K273A mutations to be responsible for the impairment of the genome encapsidation.

Finally, acetylation of K_273_, has been previously proposed to be important for different steps of the infectious cycle (78, 79). In those studies, though, the role of acetylation of K_273_ was assessed by simultaneously replacing K_264_, K_266_, and K_273_, thereby not allowing to conclude on the specific contribution of K_273_. Here, hampering acetylation but preserving the positive charge by the K273R substitution did not affect integration, indicating that the possible acetylation of K_273_ had no effect on integration. This observation is in line with the observation by Topper and co-workers that posttranslational acetylation of the integrase CTD is dispensable for viral replication (80). Altogether, the data available in the literature regarding K_240_ and K_273_ indicate that the effects we observed in this study cannot be due to any of the already known properties of the residues of the motif.

The NKNK motif is strictly conserved in natural sequences of HIV-1 group M. However, we show that various variants of the NKNK motif display levels of integration efficiency equivalent to the wt enzyme and could therefore in principle be found in the epidemics. Their absence is indicative of purifying selection occurring *in vivo*, likely exerted at a step different from integration. One possibility is the implication of IN in reverse transcription, which is, in all variants, less efficient than with the NKNK sequence. The existence of several alternative sequence arrangements, in the motif, with comparable efficiencies of integration might therefore have constituted an asset for optimizing the acquisition of additional functions, such as promoting reverse transcription.

## Materials and Methods

### Plasmids and molecular cloning

p8.91-MB was constructed by engineering one *Mlu*I and one *Bsp*EI restriction sites respectively 18 nt downstream the 5’ and 21 nt upstream the 3’ of the RT-coding sequence of the pCMVΔR8.91 (81). The insertion of the two restriction sites led to three amino acids changes in the RT (E6T, T7R and A554S). These modifications only slightly affected the efficiency of generation of puromycin-resistant clones (see below) upon transduction with the resulting viral vector (v8.91-MB) since, in three independent experiments, the number of clones obtained with p8.91-MB was 80% ± 6% of that obtained using p8.91. The p8.91-MB was employed throughout the study as positive control and was used to insert the various variants of RT and IN tested. Together with the *Sal*I site, present in the p8.91 48 bp downstream the stop codon of *pol* gene CDS, the *Mlu*I and *Bsp*EI sites define two exchangeable cassettes: one encompassing the RT coding sequence (*Mlu*I-RT-*Bsp*EI, 1680 bp) and one encompassing the IN-coding sequence (*Bsp*EI-IN-*Sal*I, 940 bp). These cassettes were used to insert the various sequences of RT and IN used in the study. The plasmid used to produce the genomic RNA of the viral vectors was a modified version of pSDY, previously described (82), hereafter called pSRP (for pSDY-nRFP-Puro). This variant was obtained by introducing two modifications to the original pSDY-dCK-Puro plasmid (82). The first one was the replacement of the sequence encoding the human deoxycytidine kinase by a cassette containing the RFP fused with the N-ter 124 amino acids of human histone H2B, which directs the RFP to the nucleus. The RFP was used to monitor the efficiency of transfection by fluorescence microscopy. The second modification was the replacement of the HIV-1 U3 sequence in the 5’ LTR by that of the U3 of the Rous sarcoma virus. For the generation of qPCR standard curves, two plasmids were constructed: one, called pJet-1LTR, for the detection of early and late reverse transcription products, was obtained by inserting the sequence encompassing the LTR and the Psi region from pSDY (82) in the pJET plasmid with the CloneJET PCR Cloning Kit (Thermo Scientific, MA, USA); the second, pGenuine2LTR, has been obtained by inserting a fragment of 290 bp corresponding to the unprocessed junction of U5/U3 (CAGT/ACTG being the sequence of the junction 5’ to 3’) into the pEX-A2 plasmid (Eurofins Genomics, Luxembourg). For the study with replication-competent viruses we used the pNL4.3 plamid (83) that was obtained from the NIH AIDS Research and Reference Reagent Program, #114 (GeneBank accession #AF324493). We replaced in this plasmid the coding sequence of NL4.3 IN CTD with those of wt and mutants INA CTD, as described in Results. Chimerical integrases between primary isolates from HIV-1 group M subtype A2 and HIV-1 group O RBF206, as well as mutant integrases, were constructed through overlap extension PCR as previously described for the envelope gene (84).

### Cells

HEK-293T cells were obtained from the American Type Culture Collection (ATCC). P4-CCR5 reporter cells are HeLa CD4+ CXCR4+ CCR5+ carrying the LacZ gene under the control of the HIV-1 LTR promoter (85). TZM-bL cells are a HeLa cell clone genetically engineered to express CD4, CXCR4, and CCR5 and containing the Tat-responsive reporter gene for the firefly luciferase under the control of the HIV-1 long terminal repeat (86). HEK-293T, P4-CCR5 and TZM-bl cells were grown in Dulbecco’s Modified Eagle’s Medium (DMEM, Thermo Fisher, MA, USA) supplemented with 10% foetal calf serum and 100 U/ml penicillin-100 mg/ml streptomycin (Thermo Fisher, MA, USA) at 37°C in 5 % CO_2_. CEM-SS cells are human T4-lymphoblastoid cells (87-89) and were grown in Roswell Park Memorial Institute medium (RPMI) supplemented with 10% foetal calf serum and 100 U/ml penicillin-100 mg/ml streptomycin (Thermo Fisher, MA, USA) at 37°C in 5 % CO_2_.

### Viral strains

The following primary isolates were used for this study: from HIV-1 group M, one from subtype A2 (GenBank accession #AF286237, named hereafter “isolate A”), one from subtype C (GenBank accession #AF286224, hereafter named “isolate C”), one from CRF02_AG (GenBank accession #MH351678), one from subtype B (isolate AiHo GenBank accession #MH351679, hereafter named isolate B); from HIV-1 group O the primary isolate RBF 206 (GenBank accession #KU168298, hereafter named “isolate O”). Isolates #AF286237, #AF286224 and #MH351678 were obtained from the NIH AIDS Research and Reference Reagent Program; isolates #MH351678, #MH351679 and #KU168298 were kindly provided by J.C. Plantier (CHU Rouen, France).

### Sequence alignments

We used 3366 HIV-1 sequences for alignment. HIV-1 group M sequences were downloaded from the Los Alamos National Laboratory (LANL) HIV sequence database and correspond to the different HIV-1 group M pure subtypes: A (249 sequences), B (2450 sequences), C (450 sequences), D (121 sequences), G (80 sequences), H (8 sequences), J (6 sequences), K (2 sequences). We also aligned 49 HIV-1 group O sequences, using 26 sequences from the LANL database and the 23 sequences obtained through collaboration with the Virology Unit associated to the French National HIV Reference Center (Pr. J.C. Plantier). Sequence alignments were performed with CLC sequence viewer 8. The sequence logo of positions 222, 240, 254 and 273 in HIV-1 group M IN was obtained with an alignment of 3366 sequences of the IN CTD using WebLogo (http://weblogo.berkeley.edu/logo.cgi).

### Generation of pseudotyped viral vectors

Pseudotyped lentiviral vectors were produced by co-transfection of HEK 293T cells with pHCMV-G (90) encoding the VSV-G envelope protein, pSRP and p8.91-MB based plasmids with the polyethylenimine method following the manufacturer’s instructions (PEI, MW 25000, linear; Polysciences, Warrington, PA, USA). HEK 293T were seeded at 5 × 10^6^ per 100-mm diameter dish and transfected 16-20h later. The medium was replaced 6h after transfection, and the vectors were recovered from the supernatant 72h later, filtered on 0.45 μm filters and the amount of p24 (CA) was quantified by ELISA (Fujirebo Europe, Belgium).

### Western blot

Western blot analysis was carried out on virions to assess the proteolytic processing of the Pr55Gag polyprotein. 1.5 mL of viral supernatant was centrifuged through 20 % sucrose, and the virion pellet was lysed in Laemmli buffer 1.5X. Viral proteins were separated on a Criterion™ TGX Strain-Free 4-15 % gradient gel (Biorad, CA, USA) (TGS: Tris Base 0,025 M/Glycine 0,192 M/SDS 0,1 %, 150V, 45 min), blotted on a PVDF membrane (TGS/Ethanol 10 %, 200 mA, 1.5h) and probed with a mouse monoclonal anti-CA antibody (NIH AIDS Reagent Program, #3537) to detect the viral capsid, the Pr55Gag unprocessed polyprotein and CA-containing proteolytic intermediates. An anti-mouse HRP-conjugated secondary antibody was used to probe the membrane previously incubated with anti-CA. Membranes were incubated with ECL reagent (Thermo Fisher, MA, USA) and WB were imaged on a Biorad Chemidoc Touch and analysed with the Biorad Image Lab software.

### Evaluation of reverse transcription by qPCR

The viral vectors were treated with 200 U/ml of Benzonase nuclease (Sigma-Aldrich, MO, USA) in the presence of 1 mM MgCl_2_ for 1h at 37°C to remove non-internalized DNA. The vectors (200 ng of p24) were then used to transduce 0.5 × 10^6^ HEK 293T cells by spinoculation for 2h at 32°C, 800 rcf, with 8 μg/mL polybrene (Sigma-Aldrich, MO, USA). After 2h, the supernatant was removed, cells were resuspended in 2 mL of DMEM and plated in 6-well plates. After 30h, cells were trypsinised and pelleted. Total DNA was extracted with UltraClean® Tissue & Cells DNA Isolation Kit (Ozyme, France). A duplex qPCR assay (see Table S2 for primers) was used to quantify early and late reverse transcription products by detecting the R-U5 and U5-Psi junctions, respectively, and another qPCR (Table S2) to normalise for the quantity of cells employed in the assay (detection of β-actin exon 6 genomic DNA; International DNA Technologies -IDT-Belgium). All primers and probes were synthesised by IDT. The qPCR assays were designed with the Taqman® hydrolysis probe technology using the IDT Primers and Probes design software (IDT), with dual quencher probes (one internal ZEN™ quencher and one 3’ Iowa Black™ FQ quencher) (Table S2). qPCRs were performed with the iTaq Universal Probes Supermix (Biorad, CA, USA) on a CFX96 (Biorad, CA, USA) thermal cycler with the following cycling conditions: initial Taq activation 3 min, 95°C followed by [denaturation 10 sec/95°C; elongation 20 sec/55°C] x 40 cycles. Standard curves and analysis were carried out with the CFX Manager (Biorad, CA, USA). DNA copy number was determined using a standard curve prepared with serial dilutions of the reference plasmids pJet-1LTR and of a known number of HEK 293T cells.

### Evaluation of integration by puromycin assay

Half a million of HEK 293T cells were transduced with a volume of viral vectors corresponding to 0.2 ng of p24, by spinoculation 2h at 32°C, 800 rcf, with 8μg/mL polybrene (Sigma-Aldrich, MO, USA). After 2h the supernatant was removed, cells were resuspended in 7 mL of DMEM and plated in 100 mm diameter plates. After 30h, puromycin was added at a final concentration of 0.6 μg/mL, clones were allowed to grow for 10 to 12 days and then counted. However, the number of clones depends on two parameters: the efficiency of integration and the amount of pre-proviral DNAs available for integration (which depends on the efficiency of reverse transcription). Therefore, we normalized the number of clones observed by the amount of viral DNA generated by reverse transcription (estimated by qPCR) to extrapolate the efficiency of integration. The percentage of integration efficiency for sample X with respect to the control C is, thus, given by (p_x_/r_x_)/(p_c_/r_c_) × 100, where r_x_ and r_c_ are the amounts of late reverse transcription products, estimated by qPCR, in sample X and in control C, respectively, and p_x_ and p_c_ the number of puromycin-resistant clones in sample X and in control C, respectively.

### Evaluation of integration by Alu qPCR

Equal amounts of total DNA extracted from transduced cells (as deduced by qPCR of β-actin exon 6 genomic DNA, see above) were used for the Alu PCR assay, as previously described (91). Two subsequent amplification were performed. The first one, 95°C for 3 min, [95°C for 30 sec, 55°C for 30 sec, 72°C for 3 min30s] x15, 72°C for 7 min, using the Alu-forward primer and the Psi reverse primer, allowed to amplify Alu-LTR fragments (Table S3). Samples were then diluted to 1:10 and 2 μL were used for the second amplification to detect the viral LTR, as described above for the detection of the R-U5 junction. The percentage of integration efficiency for sample X with respect to the control C is given by (a_x_/r_x_)/(a_c_/r_c_) × 100, where r_x_ and r_c_ are the amounts of late reverse transcription products, estimated by qPCR, in sample X and in control C, respectively, and a_x_ and a_c_ the amounts of DNA estimated by the second amplification of the Alu qPCR assay in sample X and in control C, respectively.

### Quantification of two LTR circles and of circles with perfect junctions

Non-internalised DNA was removed by treatment with Benzonase nuclease as for the qPCR assay and 0.5 × 10^6^ HEK 293T cells were transduced with a volume of viral vectors corresponding to 1 μg of p24 by spinoculation, as described above. After 30h, cells were trypsinised and pelleted. Total DNA was extracted with UltraClean® Tissue & Cells DNA Isolation Kit. Late reverse transcription products (detection of the U5-Psi junction) were quantified as described above and two qPCR assays were used to quantify the 2LTRc and the quantity of 2LTR circles with a perfect palindromic junction, with a primer overlapping the 2LTRc junction, as previously described (92). The qPCR assays were designed and the primers and probes (Table S4) synthesised as described above. qPCRs were performed as described above. Standard curves and analysis were carried out with the CFX Manager (Biorad, CA, USA). Copy numbers of the different forms of viral DNA were determined with respect to a standard curve prepared by serial dilutions of the pGenuine2LTR plasmid. The amount of 2LTRc for sample X is normalised by the total amount of the late reverse transcription products (detection of the U5-Psi junction), then it is expressed as a percentage of the amount detected for the control INA D116A (indicated with D), thus giving (2LTRc_x_/r_x_)/(2LTRc_D_/r_D_) × 100, where r_x_ and r_D_ are the amounts of late reverse transcription products, in sample X and in control D, respectively, and 2LTRc_x_ and 2LTRc_D_ the amount of 2LTR circles in sample X and in control D, respectively.

### Calculation of the efficiency of nuclear import and of 3’ processing

The efficiency of nuclear import was estimated as follows. The level of 2LTRc found with wt IN A was 0.2 with respect to that found with D116A (data from Figure 6A). The diminution observed with wt IN A with respect to D116A (which was considered to produce the maximum amount of 2LTRc and was therefore set at 1) was therefore 0.8 (given by 1 - 0.2). Since the diminution of 2LTRc is proportional to the efficiency of integration, for example a mutant integrating with an efficiency 0.3 that of wt IN A is expected to reduce the amount of 2LTRc by 0.8 × 0.3 = 0.24. The amount of 2LTRc expected for that mutant would therefore be given by 1 - 0.24 = 0.76. Similarly, a mutant integrating with a higher efficiency (for example 0.9 that of wt IN A) is expected to give 1 - (0.8 × 0.9) = 0.28 2LTRc with respect to the mutant D116A. Therefore, the formula applied to estimate the expected levels of 2LTRc with respect to D116A for sample n is given by 1 - (0.8 × a_n_) where a_n_ is the level of integration measured for sample n, relative to wt IN A (data from Figure 6A). The values of 2LTRc measured experimentally (Table 1, line 3) are then divided by the expected ones to obtain an estimate of the relative efficiency of nuclear import (Table 1, line 4).

The efficiency of 3’ processing was calculated as follows. The ratio of perfect junctions out of the total amount of 2LTRc (PJ/2LTRc) found for D116A was considered to be the maximal one and was therefore assigned a value of 1 (Table 1, line 5). The proportion of PJ/2LTRc found for wt IN A (Table 1) was 0.54 that of D116A (Table 1, line 5). The proportion by which the pool of PJ/2LTRc found with a catalytically inactive IN can be decreased by 3’ processing carried out by a fully catalytic active IN is therefore 1-0.54=0.46 (Table 1, line 6). For mutant NQNK, for example, the ratio PJ/2LTRc observed was 0.87 of D116A, which corresponds to a relative decrease of the PJ/2LTRc pool by 0.13 (Table 1, line 6). This decrease is 0.13/0.46=0.28 that observed for wt IN A (Table 1, line 7), providing an estimate of the relative efficiency of 3’ processing by this mutant with respect to wt IN A. The general formula we applied to estimate the efficiency of 3’ processing was therefore (1-r_x_)/0.46, where 1 is the proportion of PJ/2LTRc found for D116A, r_x_ is the ratio PJ/2LTRc observed for sample X and 0.46 is the decrease in PJ/2LTRc observed for wt IN A with respect to D116A. The resulting values are reported in Table 1, line 7.

### Assessment of the infectivity of replication-competent viruses

As described above, the coding sequence of NL4.3 IN CTD was replaced with those of wt and mutants INA CTD. Replication-competent viruses were produced as described above and equal amounts of viruses were used to infect cells (TZM-bL or CEM-SS) for each sample. For estimating viral replication in TZM-bL cells, 25 µL of virus dilution were added to 10^4^ cells, plated in 96 wells plates in 75 µL of culture medium. After 48h, virus replication was detected by measuring Luc reporter gene expression by removing 50 µL of culture medium from each well and adding 50 µL of Bright Glo reagent to the cells. After 2 min of incubation at room temperature to allow cell lysis, 100µL of cell lysate were transferred to 96-well black solid plates for measurements of luminescence (RLU) using a luminometer (93). For the detection of virus replication in CEM-SS cells, 0.5 ×10^6^ CEM-SS cells/5ml were infected with 1/25 virus dilution. After 5 days of culture, the percentage of infected cells were detected by intracellular p24 immuno-staining and flow cytometry analysis as previously described (94).

### Cloning, production, purification and crystallization

The C-terminal domains (IN CTD, 220-270) of integrases NL4.3, A and A K240Q/N254K studied here were cloned in the pET15b plasmid and the proteins were expressed in BL21DE3 *E. coli* cells. After transformation with the IN C-ter expressing pET15b, bacteria were inoculated at an OD_600_ of 0.1 in one litre of LB medium supplemented with 10% (w/v) sucrose. Cultures were incubated at 37°C with shaking at 220 rpm. At OD_600nm_ of 0.5, the temperature was lowered to 25°C, and shaking reduced to 190rpm, till the cells reached an OD_600nm_ of 0.8. IPTG was then added to a final concentration of 0.5 mM to induce the expression of the C-terminal domains. Cells were incubated overnight at 25°. Bacteria were then collected by centrifugation.

For protein purification, cells were resuspended in lysis buffer (25 mM HEPES pH 8, 1 M NaCl, 10 mM imidazole) in a ratio of 10 mL of buffer/gram of biomass. Roche Complete Inhibitor Cocktail tablets were added at the beginning of lysis to avoid protease degradation. Cells were lysed by sonification, for 1min/g of cells with pulse every 2 seconds at 40% amplitude at 4°C. The bacterial debris were pelleted by ultracentrifugation at 100 000xg for 1hr at 4°C. The supernatant was then loaded on a 1 mL HisTrap FF Crude column (GE Healthcare) with flow rate of 1 mL/min using the AKTA FPLC. Protein was eluted using a gradient up to 500 mM Imidazole in 10 column volumes. Protein concentration was estimated using the Nanodrop. Subsequently, the protein sample was concentrated using the Amicon Ultra 15 mL with a 3 kDa MWCO for the next purification step. A second step of purification was carried out using the S75-16/60 column (GE Healthcare) in 25 mM HEPES pH 8, 1 M NaCl. Samples were dialyzed into 25 mM HEPES pH 8, 150 mM NaCl for crystallization.

All initial crystallization conditions were determined by vapor diffusion using the TPP Labtech Mosquito Crystal. 200 nL of protein (7-4 mg/mL) was mixed with 200 nL of reservoir in 2 or 3 well of a 96 well MRC crystallization plate which was stored in the Formulatrix RockImager at 20°C. Screen included PEGS (Hampton Research), MPD, CLASSICS, NUCLEIX (Qiagen), JCSG, WIZARDS, ANION and CATION (Molecular Dimensions). Once the initial conditions were obtained, manual drops were set up in Hampton Research 24 well VDX plate to optimize crystallization conditions, and to improve crystal size and quality by mixing 1 µL protein + 1µL reservoir and equilibrating against 500 µL of reservoir at 20°C. The IN CTD NL4.3 (group M, subtype B) were obtained in a reservoir containing 0.1 M Tris pH 7, 0.8 M potassium sodium tartrate and 0.2 M lithium sulfate. For IN CTD A (subtype A2) and A K240Q/N254K the reservoir was composed of 0.1 M MES pH 6.5 and 1M sodium malonate.

### Data collection, structure solving and refinement

Data collection was performed at the Swiss Light Source (SLS, Villigen, Switzerland) on a Dectris Pilatus 2M detector. After fishing, crystals were rapidly passed through a drop of fluorinated oil (Fomblin® Y LVAC 14/6, average MW 2,500 from Sigma Aldrich) to prevent ice formation and directly frozen on the beamline in the nitrogen stream at 100 K. X-ray diffraction images were indexed and scaled with XDS (95, 96). The structures were solved by molecular replacement using PHASER (97) in the PHENIX (98) program suite using the NMR HIV-1 C-ter structure (1QMC) (99) as a search model for the IN CTD NL4.3 structure, which was used subsequently as a search model to solve the A and A K240Q/N254K structures. The structure was then built using the AUTOBUILD program (100, 101) followed by several cycles of refinement using PHENIX.REFINE (102) and manual rebuilding with COOT (103). Structure based sequence alignment was performed using PROMALS3D (104). Structures superposition and Root Mean Square Deviations (RMSD) calculations have been performed using secondary structure matching (SSM), superpose program (105) embedded in COOT (103) and in the CCP4 program suite (106). The sequence alignment representation has been generated by ESPript (107). Surface potential was calculated using the DELPHI web server (108) and visualized with CHIMERA (109). Data collection and refinement statistics are summarized in Table S5. Crystallographic structures were deposited in PDB under the identification numbers 6T6E (HIV-1 Cter, PNL4.3), 6T6I (HIV-1 Cter, subtype A2) and 6T6J (HIV-1 Cter, subtype A2, mutant N254K-K240Q).

### Analysis of the surface electrostatic potential of retroviral CTDs

The structures and the sequences of the C-terminal domains have been extracted from: HIV-1 A2, PDB 6T6I (this publication); HIV-1 PNL4.3, PDB 6T6E (this publication); SIV, PBD 1C6V (48); MVV, PDB 5LLJ (49); RSV, PDB 1C0M (50); MMTV, PDB 5D7U (51); MMLV, PDB 2M9U (52); PFV, PDB 4E7I (53). The structure based sequence alignment has been performed using PROMALS3D (104). Structures superposition and rRoot Mean Square Deviations (RMSD) calculations have been performed using secondary structure matching (SSM), superpose program (105) embedded in COOT (103). The sequence alignment representation has been generated by ESPript (107). The surface electrostatic potential was calculated using the DELPHI web server (108) and visualized with CHIMERA (109).

### Statistical tests

All statistical analyses were performed on at least three independent experiments (transfection and transduction) using Prism 6 (GraphPad). For all functional tests, the values obtained for the chimeras were normalized using the values obtained for parental integrase A. Student tests were used to evaluate whether the normalized mean values obtained with the chimeric and mutant integrases were significantly different from that obtained with the parental strain, and/or between them. For confocal microscopy, unpaired t test was used for statistical analyses.

## Acknowledgements

The authors are grateful to Pr. J.C. Plantier for providing HIV-1 strains of subtype B, CRF02 and group O, to C. Elefante for the construction of the the p8.91-MB plasmid, to J. Batisse for providing control reagents, and to M. Lavigne, B. Maillot and S. Marzi for helpful discussions. The authors wish to thank R. Drillien (IGBMC) for suggestions about the manuscript. The authors thank V. Olieric and the staff of the Swiss Light Source synchrotron for help with data collection. The authors acknowledge the support and the use of resources of the French Infrastructure for Integrated Structural Biology FRISBI ANR-10-INBS-05 and of Instruct-ERIC.

